# Structural Determinants of Tendon Function During Development and Their Sensitivity to Mechanical Stimulation

**DOI:** 10.1101/2023.08.25.554314

**Authors:** Benjamin E. Peterson, Maria L. Canonicco Castro, Helen McCarthy, Niamh Buckley, Nicholas Dunne, Rebecca Rolfe, Paula Murphy, Spencer E. Szczesny

## Abstract

The load-bearing capabilities of tendon are acquired during neonatal stages of development, characterized by an abrupt increase in multiscale mechanical properties. While prior work has identified numerous changes within the collagenous structure during these developmental periods, the primary structural elements that give rise to this abrupt mechanical functionality, and their mechanobiological sensitivity, remains unclear. To address this gap in knowledge, we leveraged a combination of ultrastructural imaging, biochemical/thermodynamic assays, multiscale mechanical testing, and shear lag modeling to probe the dynamic structure-function relationships and establish their sensitivity to mechanical stimulation during tenogenesis. Mechanical testing and modeling suggested that the rapid increase in multiscale mechanics can be explained by a increasing fibril length and intrafibrillar crosslinking. To test this, we inhibited collagen crosslinking during development and observed a drastic reduction in multiscale mechanical capabilities that was explained by a reduction in both fibril modulus and length. Using muscle paralysis to investigate mechanosensitivity, we observed a significantly impaired multiscale mechanical response despite small changes in fibril diameter and fibril area fraction. While there was no change in crosslinking density, there was a decrease in thermal stability with flaccid paralysis, and our shear-lag model suggested that flaccid paralysis produces a reduction in fibril length and intrafibrillar crosslinking. Together, these data suggest that both intrafibrillar crosslink formation and fibril elongation are critical to the formation of load-bearing capabilities in tenogenesis and are sensitive to musculoskeletal activity. These findings provide critical insights into the biological mechanisms that give rise to load-bearing soft tissue.

## 1. Introduction

Tendon is a highly collagenous connective tissue that transmits force between muscle and bone enabling locomotion. The characteristic high load-bearing capabilities of tendon are acquired during embryonic and neonatal stages of development^1–3^ facilitated by the coordinated deposition and organization of fibrillar collagens^4–6^. While the timing may vary, this developmental process is highly conserved across species and results in the formation of a complex hierarchical structure that spans length scales across multiple orders-of-magnitude^7^. While prior work has provided significant insight into the dynamic structural and mechanical changes that occur during tendon development, the field lacks a comprehensive understanding of the structure-function relationships in developing tendons and the relative importance of each structural change to the emergent mechanical properties. Additionally, while mechanical stimulation is known to be important for proper tendon development^8–11^, it is still unclear which structural elements are mechanosensitive and require embryonic muscle activity. A greater understanding of the structure-function relationships during tendon development is required to elucidate the complex biological and mechanical cues required for tendon formation and may provide key insights to improve tissue engineering and regenerative medicine strategies.

Tendon development is initiated by the expansion and linearization of tendon fibroblasts (tenocytes), which ultimately define the longitudinal axis of the tendon^12^. Over time, soluble procollagen molecules (primarily collagen I) are secreted into aligned extracellular channels, which then undergo a series of post-depositional modifications to enable collagen fibril assembly. This process, termed fibrillogenesis, is tightly regulated via interactions with other fibrillar collagens (V and XI), matrix metalloproteinases, and proteoglycans at the cell surface to produce short intermediate fibrils approximately ∼10 um in length and ∼20 nm in diameter^2,5^ Over time, these short intermediate collagen fibrils begin to accumulate within the extracellular matrix, marked by a substantial reduction in cellular volume fraction^2,13^. This initial phase is highly conserved across animals, initiating at embryonic day 14.5 (E14.5) in mice^12^ and E10 in chicks^14^. During this period, fibrils undergo small increases in their length and diameter; however, there are minimal changes to tendon gross tensile properties^1,3,13^.

At later developmental stages there is an abrupt reorganization of the extracellular matrix as fibrils undergo a coordinated series of longitudinal and lateral fusion events, respectively increasing their length and diameter^5,14^. This process begins around E16 in chicks and post-natal day 10 in mice and corresponds with a rapid increase in macroscale mechanical capabilities, marking the transition towards functional load-bearing capabilities^1,2,10^. Indeed, recent work from our lab has demonstrated an increase in multiscale mechanical capabilities during late stages of embryonic chick development (E16 - E20)^10^. More specifically, we observed an increase in interfibrillar load transmission with maturation, consistent with a progressive increase in collagen fibril lengths. We further demonstrated that the induction of flaccid muscle paralysis prior to this structural transformation can impair the maturation in multiscale mechanical capabilities, highlighting the importance of external mechanical stimuli in development. Despite these data, the relative importance of fibril lengthening on tendon mechanics compared to other simultaneous structural changes that occur during development, and their sensitivity to muscle stimulation, is still unclear. Recent work has suggested that intrafibrillar crosslinking, mediated by lysyl oxidase (LOX) activity, may play a key role in enabling load-bearing capabilities during tendon development^15,16^. Indeed, the presence of mature intrafibrillar crosslinks, hydroxylysl-pyridinoline (HP) or lysyl-pyridinoline (LP), increases with tendon maturation and corresponds with an increase in tissue stiffness^17^. Additionally, developmental muscle paralysis resulted in a reduction in LOX activity and compressive modulus, suggesting that impaired intrafibrillar crosslink formation may be contributing to the structure-function phenotype following paralysis^11^. Furthermore, *in vitro* constructs have demonstrated irregular fibril morphology following LOX inhibition^18^. Together, these data suggest that LOX is a mediator of fibril fusion and is modulated via muscle activity during development. However, a more comprehensive assessment of the structure-function changes that occur during late stages of embryonic development and their respective sensitivity to muscle stimulation is required to elucidate their contributions to the production of robust load-bearing tendons.

Therefore, the objective of this study was to conduct a comprehensive characterization of the dynamic structure-function relationships throughout the late stages of tendon development and to evaluate how mechanical stimulation affects the establishment of these relationships. To accomplish this objective, we conducted a combination of multiscale mechanical testing, shear-lag modeling, biochemical/thermodynamic assays, and ultrastructural imaging of embryonic chick tendons. We hypothesized that fibril elongation and collagen crosslinking will be the primary structural changes required for the proper maturation of tendon function and that the absence of muscle activity during development will impair both crosslink maturation and fibril elongation. These findings will provide critical insight into the structural mechanisms that drive the formation of mature tendon mechanical capabilities and the role of mechanobiology during tendon development.

## 2. Methods

### 2.1 Sample Collection and Treatments

Fertilized chicken eggs (White Leghorn) were acquired at embryonic day 0 (E0) from the Penn State Poultry Education and Research Center and incubated in at 37.7°C. At E3, eggs were windowed following standard protocols in the field^10^.

At E13, paralysis treatments were initiated via the dropwise administration of solutions to the chorioallantoic membrane (CAM). Rigid paralysis was induced with a 100 µL of 0.2% decamethonium bromide (DMB, Sigma Aldrich) in HBSS + 1% antibiotic-antimycotic (AA, Thermo Fisher Scientific). Flaccid paralysis was induced with 100 µL of 0.2% pancuronium bromide (PB, MP Biomedicals) in HBSS + 1% AA. Both modes of paralysis were maintained with subsequent treatments from E14 – E19 with 50 µL of their respective solutions. Vehicle controls were treated with HBSS and AA alone at matching volumes.

Lysyl oxidase (LOX) mediated crosslink formation was inhibited via the treatment of β-aminopropionitrile (BAPN, Thermo Scientific) to the chorioallantoic membrane. BAPN irreversibly blocks LOX and LOXL-4 activity, impairing aldehyde formation at telopeptide lysine or hydroxylysine residues, impeding downstream crosslink formation ^19–21^. Embryos were treated at E15 with 400 uL of a 5 mg/mL solution of BAPN. Solutions were prepared with sterile HBSS + 1% AA. Vehicle controls were treated with HBSS and AA alone.

Embryonic motility was assessed during development following paralysis or crosslink inhibition treatments, as outlined in prior work^10,22^. Briefly, embryos were visually examined and any discernable movements (i.e., kicking or twisting) were counted over a 2-minute interval. Embryos were selected at random (n = 5 - 6) and motility counts (events / 2 mins) were averaged to get a representative value for each condition. Motility was examined daily from the initiation of treatment through E19.

Treated embryos were sacrificed at E20 via a 15-minute exposure to −20°C immediately followed by decapitation. To assess any changes with development, windowed embryos were sacrificed at E16, E18, and E20, but otherwise saw no treatment. All embryos were staged using Hamburger and Hamilton (HH) criteria^23^ to ensure proper developmental comparisons at E16 (HH42), E18 (HH44), or E20 (HH46). Embryo hindlimbs were isolated, wrapped in gauze, hydrated in phosphate buffered saline (PBS), and stored at −20°C.

### 2.2 Multiscale Mechanical Testing Sample Preparation & Testing

Multiscale mechanical testing was conducted as previously described in literature^10,24,25^ to assess changes in micro/macroscale mechanical behavior following paralysis and crosslink inhibition. Briefly, the flexor digitiform brevis (FDB) digit II tendon was isolated from the hindlimb, stained with 5 ng/mL dichlorotriazinylaminofluorescein (5-DTAF, Invitrogen), and mounted in custom grips at a 10 mm gauge length. Cyanocacrylate (Loctite 713) was used to ‘pot’ the tissue ends to minimize strain concentrations at the grips. Grips were then placed in a custom-built microtensile device that sits atop a confocal microscope (Nikon A1R HD). This device includes an integrated water bath filled with PBS to maintain tissue hydration during the entirety of the testing period. A pre-load of 0.1 g was applied and sets of photobleached lines (PBL) (4 lines, 80 µm apart, 3 µm wide) were bleached at the sample center and ± 1.5 mm from the center. Reference z-stack images were collected at all PBL sites and 3D surface profiles were utilized to measure the cross-sectional area assuming an elliptical cross-section.

Samples were loaded in 2% strain increments at rate of 10%/min, followed by a 20-minute stress relaxation period. A prolonged relaxation period was allowed to ensure that the tissue stress had stabilized and that the tissue surface was not actively deforming prior to image acquisition. At the end of the relaxation period, z-stack images were acquired at all PBL locations to optically track displacement behavior and the physical confocal stage position was recorded. This process was iteratively repeated until full or partial sample rupture, defined as a reduction in tissue stress from the prior increment.

#### Image Processing & Data Analysis

Multiscale mechanical data was processed as previously published in literature using a custom MATLAB (MathWorks) script^26^. In brief, a Sobel detection strategy was utilized to create a 2D projection of the curved tendon surface. The center of the photobleached line was then defined along the tissue width as the minimum local pixel intensity. Local fibril strains were then calculated by measuring the change in distance between the PBLs at each pixel location along the tissue width following loading. The measured fibril strains were then averaged within a given set of PBLs, and then across the three PBL sites, to generate a single representative value for fibril strain per loading increment. Note that our measurements of fibril strains are representative values as our imaging resolution (0.66 μm/px) cannot directly deconvolve individual fibril strains due to their smaller diameter (∼60 nm)^2^ and are unlikely to exist perfectly longitudinal to the imaging plane.

To account for gripping artifacts, the macroscale bulk tissue strains were optically calculated utilizing the displacement of the peripheral (±1.5 mm) PBL sites. More specifically, the custom MATLAB scripts were utilized to determine the centroid of the PBLs within the captured z-stack. These data along with the physical stage positions were used to measure the applied macroscale strains at each loading increment.

The fibril:tissue strain ratio was calculated at each strain increment by dividing the average fibril strain over the macroscale tissue strain. Prior work from our lab has demonstrated that a fibril:tissue strain ratio less than one is representative of interfibrillar shear between discontinuous fibrils. The fibril:tissue strain ratio will approach one as more strain is transmitted to the fibrils and the interfibrillar shear decreases. The equilibrium stress and modulus was calculated by first applying a moving average (k = 30 data points) to smooth the load cell data. The equilibrium stress for each strain increment was calculated by averaging the last 30 seconds of the stress-relaxation period and plotted against the macroscale tissue strain. The equilibrium modulus was calculated by fitting a line through the equilibrium stress-strain curve prior to sample failure. Note that samples that failed early may only be fit across ∼3 strain increments.

### 2.3 Ultrastructural Imaging

#### Sample Preparations & Imaging

Untreated windowed chick embryos were sacrificed at E16, E18, and E20 via decapitation and hindlimbs were isolated. The hindlimbs had their skin and Achilles tendon removed so the flexor tendons had direct contact with the fixative solution. Limbs were immersed in a freshly prepared fixative solution of 2% paraformaldehyde/2.5% glutaraldehyde with 2 mM calcium chloride in 0.15M cacodylate buffer (pH 7.4). After 3 days, the samples were removed from the fixative solution, FDB digit II tendons isolated, and stained and resin embedded *en bloc* utilizing standard protocols^27^. Identical sample preparation was conducted to assess the ultrastructural changes following muscle paralysis (DMB & PB) and cross-link inhibition (BAPN) at the E20 timepoint. All samples were prepared in biological replicates (n = 2 per condition). While this sample size was not sufficient to make statistical comparisons across groups, it was useful to obtain estimates of the fibril volume fraction for shear-lag modeling.

Transverse cross-sections were scanned in series at 2.5 kV under low-vacuum pressure (35 - 40 Pa) using an Apreo VolumeScope scanning electron microscope (SEM) (Thermo Fisher Scientific, Waltham, MA). All samples were imaged with a 4096 x 4096 px field of view at a resolution of 2 nm/px. Six representative images were captured for each biological replicate to account for regional variance in fibril characteristics.

#### Image Analysis

A deep learning model was used automate the fibril selection process, enabling us to screen more regions across the sample and increase the quantity of fibrils counted. Notably, all images had variance in stain/image intensity, fibril quality, and packing density which made it difficult to produce a single robust deep learning model for fibril segmentation. Thus, a unique deep learning model was produced for each sample condition, based on manually segmented regions, to tailor the fibril identification to the quality of each image set. In an effort minimize user bias in the production of the training sets, all model training was conducted prior to any quantitative analysis.

Raw SEM images underwent a series of initial functions to enhance image quality prior to training the deep learning models. First, raw images underwent a Gaussian blur filter (3 px) to eliminate noise. The resulting image stack then underwent a Fast-Fourier bandpass filter to remove large structures (i.e., shading correction, down to 40 px) and small structures (i.e., smoothing, up to 3 px) by Gaussian filtering in the Fourier space. This resulted in an increase in edge sharpness and amplifying the detection of individual fibril boundaries.

En mass fibril identification was then conducted via a deep learning strategy using Dragonfly (Version 2022.1; Object Research Systems). A unique segmentation model was trained for each distinctive timepoint/treatment condition. Each model was trained by manual segmentation of ∼40% of the field of view from two technical replicate images. Regions of interest (ROI) were selected in an effort to capture variability in the image and enhance fibril identification. Deep learning models were trained using a 2D U-Net architecture^28^ and the supplied training subsets. Additional data augmentations enabled using system defaults. These data augmentations increase training subsets by inducing artificial image transformation (i.e., shear, rotation, intensity variance, etc), which enable the model to train across additional data sets. Following training, models were applied to the entire data sets and manually checked to ensure segmentation quality. Notably, intracellular components that may appear fibril-like when transversely sectioned (such as mitochondria) were manually removed to prevent incorrect fibril labeling within the cells.

The resulting fibril masks were binarized and exported to FIJI for particle analysis. More specifically, masks were first watershed to separate adjacent touching fibrils. The particle analysis function used to measure particles with an area greater than 500 nm^2^ and a circularity between 0.3 – 1.0. These parameters were selected to filter out small particles produced by the mask water-shedding based on previously reported fibril size metrics^2,12^. Fibril diameter was calculated from the fibril area assuming a circular cross-section. Measurements from were pooled to produce a single histogram of the fibril diameter distribution for each developmental/treatment condition. The fibril area fraction was calculated for each image by diving the cumulative fibril area by the total area of the fields-of-view.

### 2.4 Shear-Lag Modeling

#### Model Fitting

An established elastoplastic shear lag model that accurately describes the multiscale mechanics of tendon fascicles^29^ was used to fit the data from embryonic chick tendons treated with DMB, PB, and BAPN. Additionally, the model was used to fit prior experimental data collected from normally developing E16, E18, and E20 tissue^10^. The model contains 8 parameters: fibril radius, fibril volume fraction, fibril modulus, fibril length, average fibril uncrimping stretch, standard deviation of fibril uncrimping stretch, elastic interfibrillar shear modulus, and plastic limit of interfibrillar shear stress. The fibril radius and fibril volume fraction were assigned for each sample group based on the average values obtained from the ultrastructural imaging measurements. For each set of experiments (i.e., development, paralysis, crosslink inhibition), the multiscale mechanics of the E20 control tissue was fit by the elastoplastic model using the lsqcurvefit function in MATLAB R2023a (MathWorks) and the initial value estimates in **Supplemental Table 1**. First, the macroscale stress-strain curve was fit by itself to fine tune the initial guesses of the uncrimping model parameters. Then the full multiscale mechanical data were fit simultaneously while bounding the uncrimping model parameters to within ± 20% of the pre-fit values to ensure that the low-stress nonlinear region of the stress response was properly fit. Note that the macroscale stress-strain data and fibril:tissue strain ratio data were normalized so that the curve-fitting function weighted both datasets equally. Because the plastic limit of the interfibrillar shear stress is inversely proportional to the fibril length^26^, it was not possible to determine unique values for these two parameters from the model fits. Therefore, we assumed a fixed value of 2 mm for the collagen fibril length in the E20 control samples, which is based on data from mouse tendon development and prior estimates of collagen fibril lengths^12,24^. For all the other samples harvested at earlier timepoints or exposed to a chemical treatment, we fixed the plastic limit of interfibrillar shear stress to the value determined from the fit of the respective E20 data. This enabled us to determine the predicted change in fibril length necessary to fit the multiscale data at earlier timepoints (i.e., E16 and E18), with paralysis (i.e., DMB and PB), and with crosslink inhibition (i.e., BAPN). All the other model parameters were allowed to vary to provide the best fit of the data and had the same initial guesses. The only other difference was that the initial guess for the fibril length for these samples was determined by inverting the equation for the plastic limit of interfibrillar shear stress (**Supplemental Table 1**).

#### Model Simulations

The elastoplastic model was also used to predict what would happen to the tendon multiscale mechanics if fibril elongation, collagen crosslinking, or changes to fibril diameter and volume fraction was inhibited during development. To accomplish this, stress-strain and fibril:tissue strain ratio curves were generated using the model parameters determined from fitting the normally developed E20 tendons. However, to mimic a failure in fibril elongation, the fibril length was set to the value determined by fitting the E16 tendons. Alternatively, to mimic inhibition of crosslink formation, the fibril modulus was set to the value determined by fitting the E16 tendons. Similarly, to mimic inhibition of lateral fibrillar growth, either the average fibril diameter or fibril volume fraction was set to the value determined from ultrastructural imaging of E16 tendons. The plots of each condition were graphed together with the normal E20 behavior for visual comparison.

### 2.5 Crosslinking Analysis

#### Differential Scanning Calorimetry

Crosslink formation alters the thermodynamic transition from a highly ordered helix to a denatured arrangement of randomly ordered α-chains^30^. This change in thermodynamic transition was assessed using differential scanning calorimetry (DSC) to provide a quantitative assessment of functional collagen crosslinking^30,31^ in tendons during physiological development and under paralysis/BAPN treatments. To increase tissue mass, both the FDB & FDL digit II tendons were isolated from each avian hindlimb and pooled. Samples were blotted on lens paper to remove excess moisture and weighted. Samples were placed within an aluminum DSC pan (TA Instruments, T210707) and gently pressed into the bottom of the pan to maximize the tissue contact area. The lid (TA Instruments, T210416) was added, and the pan was hermetically sealed. Calorimetry was performed using a DSC Q2000 (TA Instruments, New Castle, DE) accurate to 0.1°C. Samples were allowed to equilibrate at 30°C and then ramped to 70 - 75°C at a rate of 3°C/min ^31^. All tests were conducted using a sealed empty pan as an internal reference.

DSC endotherms were analyzed using TA Universal Analysis software (Version 4.5A, TA Instruments) to collect the peak denaturation temperature (T_peak_) and specific enthalpy of denaturation (Δh) for each distinctive endotherm present (**Fig. 1**). Utilizing the analysis software, a tangential flat line was drawn across each endotherm peak. The T_peak_ was calculated as the minima of the endotherm curve, describing the peak change in heat flow and the point at which ∼50% of the protein has denatured. The specific enthalpy of denaturation for each distinctive peak was calculated by the TA Universal Analysis software as the area between the drawn tangent line and the endotherm curve (**Fig. 1**). The energy for denaturation at the lower temperature was then calculated relative to the overall reaction (Peak 1 Δh / Total Δh) to assess the amount of collagen with a diminished crosslinking state, normalized to the tissue mass and collagen content.

**Figure 1:**
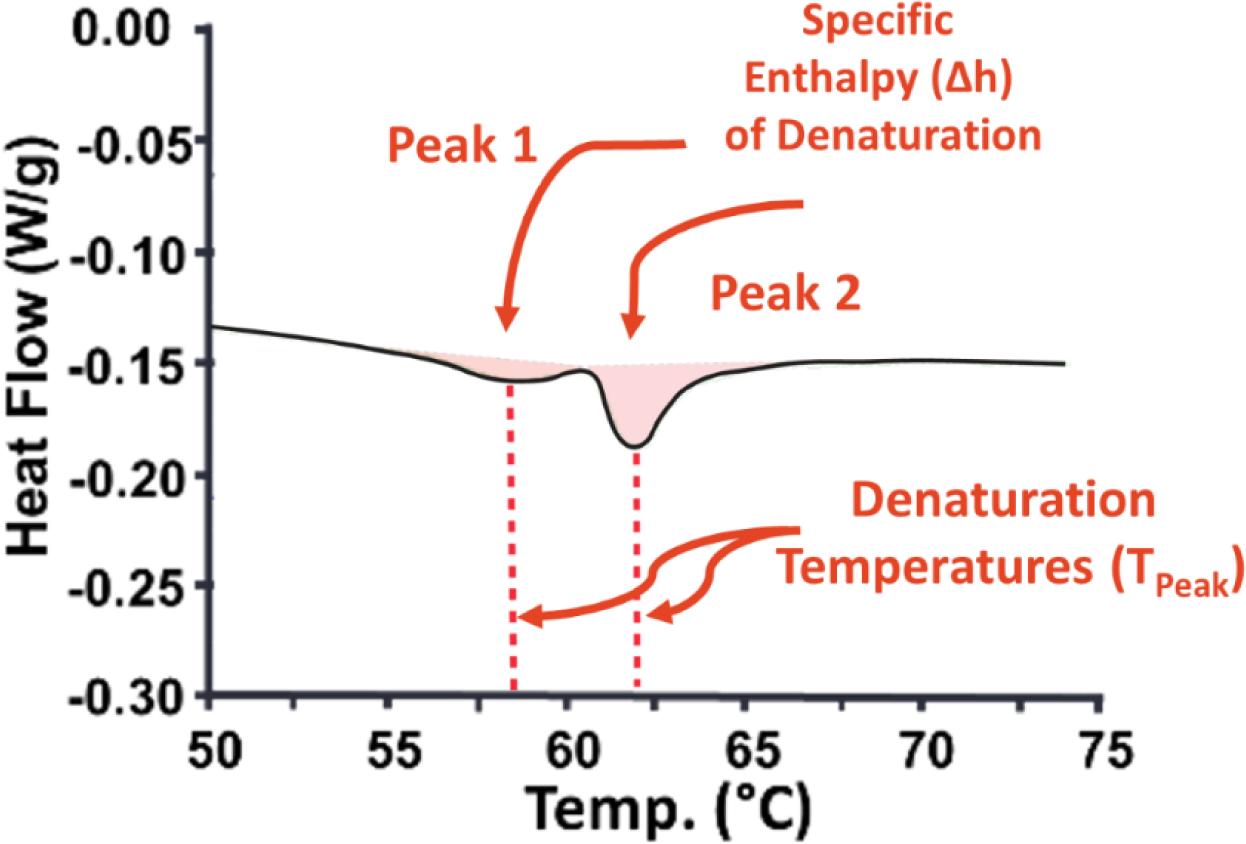
Representative differential scanning calorimetry endotherm. Multiple peaks represent unique populations of protein with distinctive denaturation behavior. T_peak_ describes the temperature at which *protein* denaturation is maximal. Area under the curve *(light red)* represents the specific enthalpy of denaturation (Δh).

#### Biochemical Assays

Mature HP and LP crosslink populations were determined utilizing commercially available kits, MicroVue Serum PYD (Quidel Crop., Kit #8019) and MicroVue DPD (Quidel Corp., Kit #8007), respectively. To ensure adequate tissue mass, both the FDB/FDL digit II tendon were isolated from both avian hindlimbs and pooled. Dissected tendons samples were lyophilized for 5 days, and the dry weight was measured. For all assays, an average dry weight of 5.17 mg (± 1.78 mg) was observed. Following lyophilization, samples were hydrolyzed in 6 N HCl for 24 h at 110°C. Residual HCl was evaporated off, reconstituted at 5 mg dry mass/mL in double distilled water, and then stored at −20°C until testing. Tendons samples were diluted to ensure measurements were within the sensitive range of the standard curve^32^. For HP analysis, samples were diluted 25-fold, while samples for the LP analysis were kept at the original 5 mg/mL concentration. Samples were otherwise prepared following manufacturer’s specifications. Tests were conducted in biological triplicate and technical duplicate.

### 2.6 Statistical Analysis

For mechanical characterization, statistical analysis was conducted using SAS (9.4) and GraphPad Prism (8.3.0). Differences in the fibril:tissue strain ratio were evaluated using a linear mixed model that takes into consideration the correlated strain increments per specimen as a covariate. A one-way ANOVA with a Tukey’s post-hoc test corrected for multiple comparisons was conducted to evaluate differences in ultimate tensile stress, modulus, fibril area, and crosslink populations as a function of development. A post-hoc Dunnett’s test corrected for multiple comparisons was likewise used to evaluate measurement differences as a function of BAPN or paralysis treatments and to evaluate differences in thermal properties in comparison to the E20 timepoint. Differences in fibril distributions were compared using Kolmogorov-Smirnov tests. The quality of the model fits was quantified by the Nash-Sutcliffe efficiency (R^2^), which is analogous to the coefficient of determination except that values less than 0 are possible and indicative that the model performs worse than a horizontal line located at the mean experimental value^33^.

## 3 Results

### 3.1 Normal Tendon Development

#### Ultrastructural Changes with Development

During development, we observed small but statistically significant changes in the fibril diameters with development. Specifically, at E18 there was an increase in the number of larger diameter fibrils that persisted at E20 (p < 0.001) (**Fig. 2**). Similarly, with increased maturation, there was trend toward an increase in fibril area fraction (p = 0.07).

**Figure 2:**
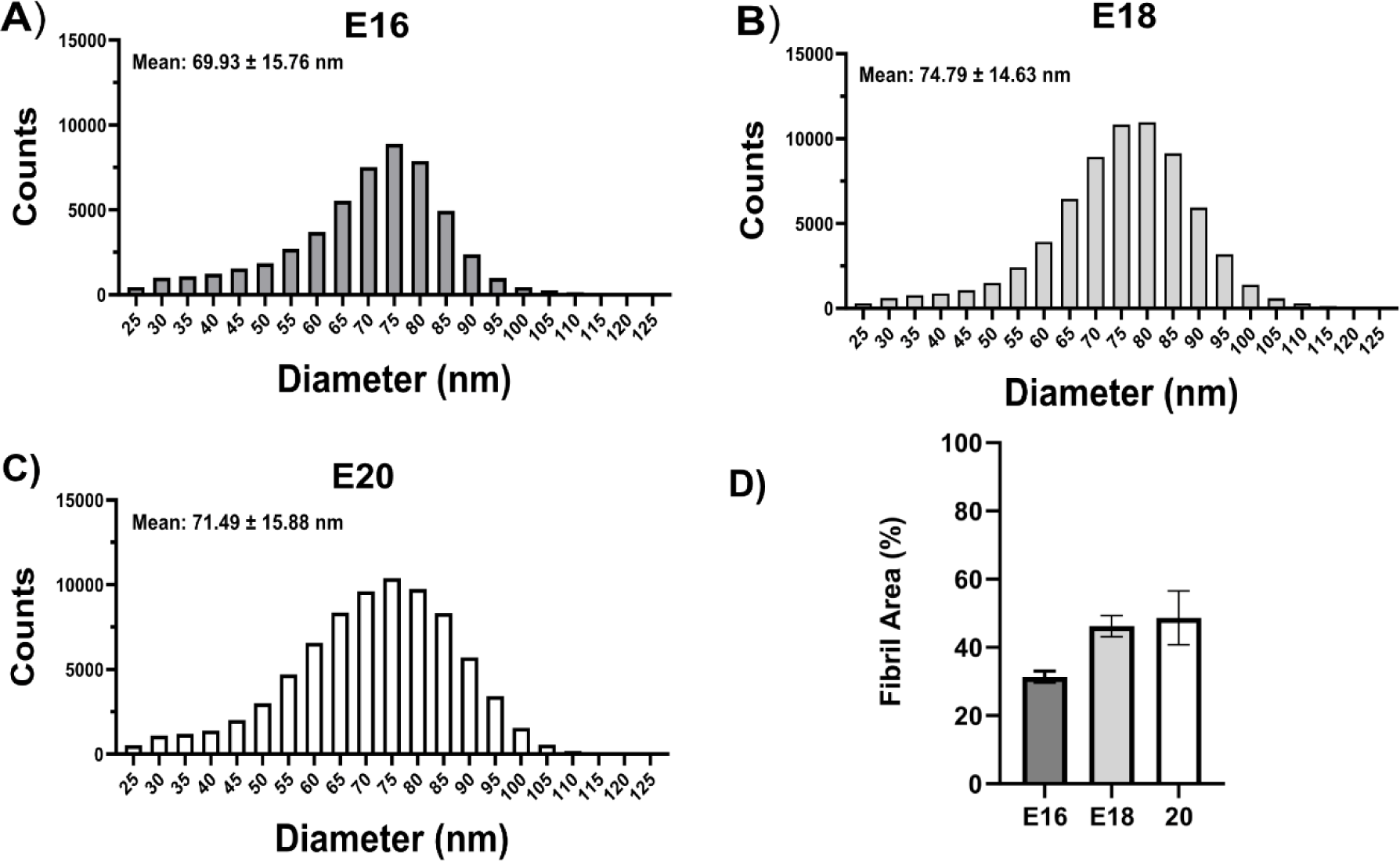
Ultrastructural changes during development. **A - C)** Fibril diameter plots for embryonic days 16, 18 and 20. For all developmental timepoints there is a unimodal distribution of fibrils. **D)** There was a trend toward an increase in fibril area with increased maturation (p = 0.07). Each timepoint consisted of two biologicals replicates (n = 2) and six representative regions of interest.

#### Multiscale Mechanical Response and Shear-Lag Modeling

In general, the elastoplastic shear lag model did a good job of fitting the prior data of embryonic tendon multiscale mechanics at all developmental ages (**Fig. 3**). The R^2^ values for all of the stress-strain and fibril:tissue strain ratio curves were above 0.98 and 0.75, respectively. The values for all the model parameters varied substantially across developmental age (**Table 1**). Specifically, the fibril modulus, fibril length, average uncrimping stretch, and standard deviation of uncrimping all increased from E16 to E20. Conversely, the elastic interfibrillar shear modulus decreased by several orders of magnitude with increasing age.

**Figure 3:**
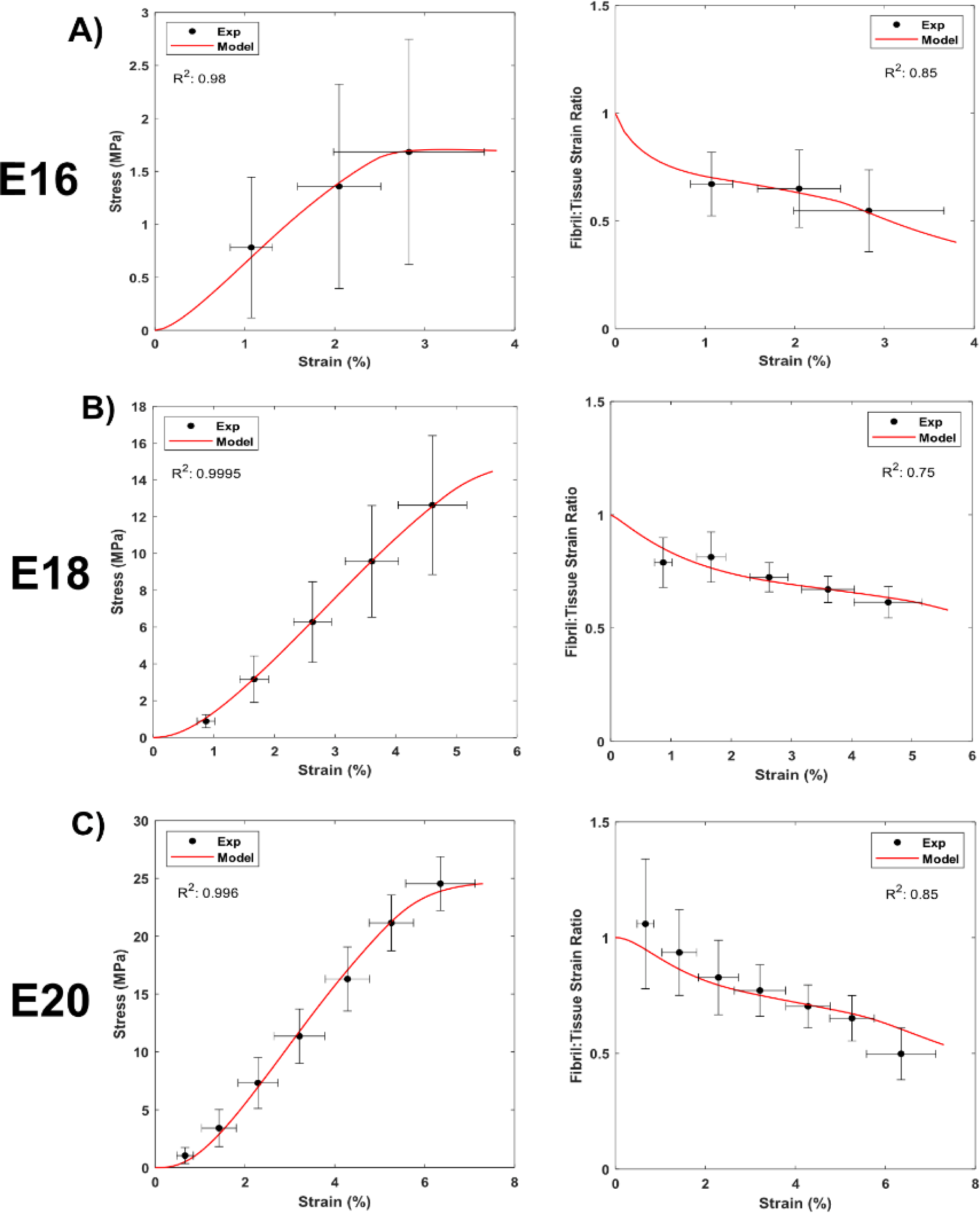
Shear lag model fit of multiscale mechanical behavior during embryonic development. The model successfully fit the macroscale mechanics and the fibril:tissue strain ratio during late stages of embryonic chick tendon development. Note that the experimental data were originally reported in a prior study^10^.

**Table1:** Fit shear-lag model parameters during embryonic development.

|  | *Fibril Modulus (MPa)* | *Fibril Length (mm)* | *Avg. Uncrimping Stretch* | *Std. Dev. Uncrimping Stretch* | *Shear Modulus (kPa)* | *Plastic Limit (kPa)* |
| --- | --- | --- | --- | --- | --- | --- |
| **E20** | 1775 | 2 (fixed) | 1.0104 | 0.0071 | 0.0055 | 1.8 |
| **E18** | 1306 | 1.36 | 1.0083 | 0.0075 | 0.0071 | 1.8 (fixed) |
| **E16** | 419 | 0.22 | 1.0023 | 0.0026 | 0.2 | 1.8 (fixed) |

#### Crosslinking Maturation

Biochemical ELISAs (**Fig. 4A-C**) demonstrated a non-significant but trending increase in mature HP+LP crosslinks with maturation (n = 3, p = 0.096). The DSC data exhibited two distinctive endotherm peaks across all developmental timepoints (**Fig. 4D**). With development, there was an increase in total specific enthalpy of denaturation and peak denaturation temperatures in comparison to the E20 timepoints (**Supplemental Table 2**). The Peak 1 enthalpy / total enthalpy ratio was used to assess the amount of more easily denatured protein proportionate to the entire collagen population. A significant reduction in Peak 1/ total enthalpy was observed between the E18 and E20 timepoints (p < 0.05), which is suggestive of increased functional crosslinking.

**Figure 4:**
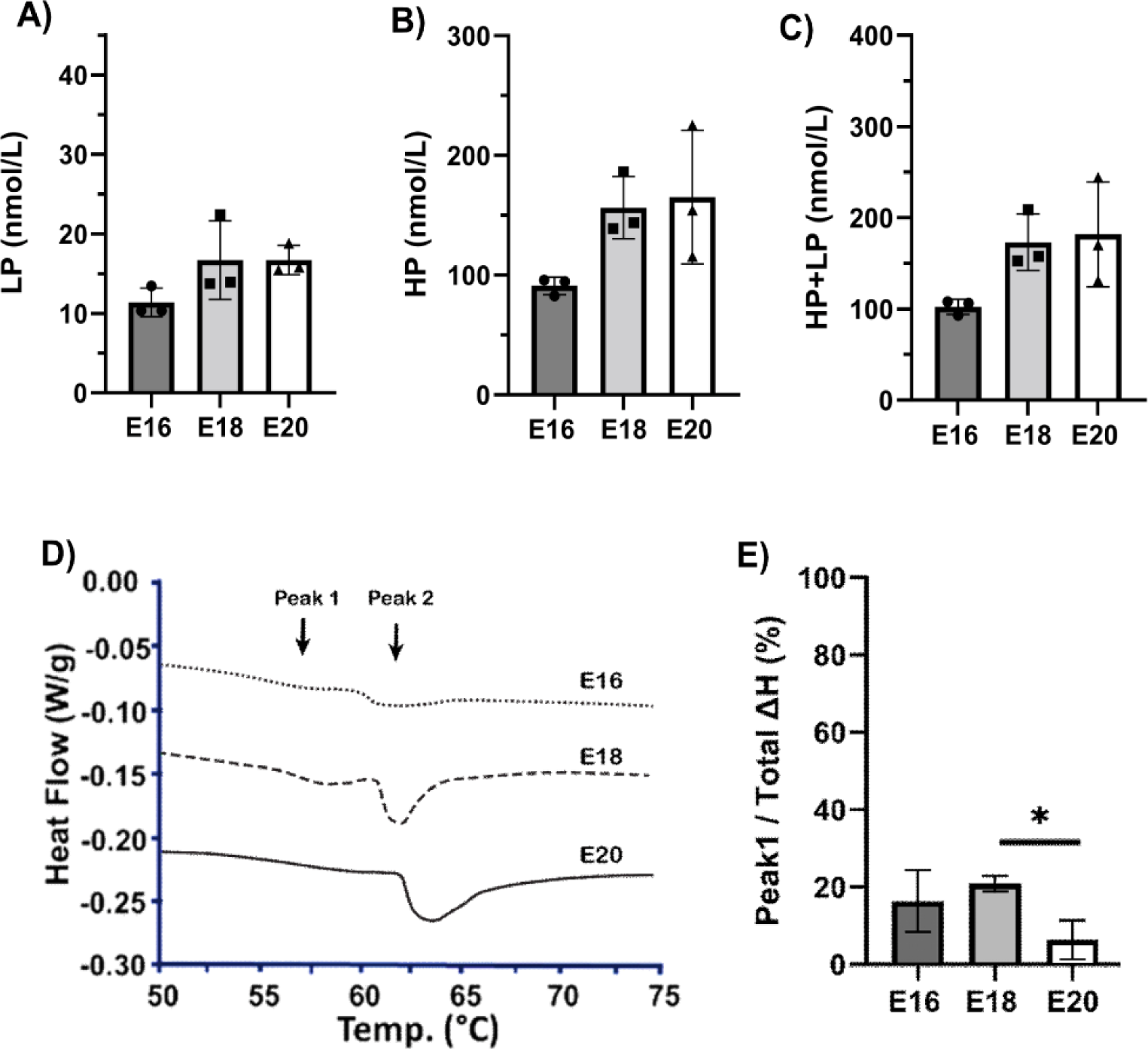
Changes in crosslinking with development. Biochemical quantification of mature A) LP and B) HP crosslinks as a function of development. C) Over development (n = 3), there is a trend toward an increase in mature crosslinks (HP +LP) (p = 0.096). D) Representative endotherms at embryonic day 16, 18, and 20. Arrows mark the two peaks of enthalpy absorption. Endotherms shifted on the y-axis to avoid overlap. E) Peak 1 enthalpy/ total enthalpy plot. Data represented as mean ± standard deviation. * p< 0.05.

### 3.2 Individual Contributions of Structural Changes to Tendon Mechanics During Development

To estimate the independent contributions of fibril elongation, collagen crosslinking, fibril diameter, and fibril volume fraction to the maturation of embryonic tendon multiscale mechanics, the elastoplastic shear lag model was used to simulate the behavior of E20 chick tendons that had failed to elongate their fibrils, produce more crosslinks (i.e., increase the fibril modulus), increase fibril diameters, or increase fibril volume fraction compared to E16 tendons. These simulations suggest that both fibril elongation and crosslinking are critical to normal tendon development (**Fig. 5A-B**). Specifically, prevention of fibril elongation dramatically reduces the stress response of the tissue as well as the fibril:tissue strain ratio, which is consistent with less strain being transmitted to the fibrils due to increased interfibrillar sliding. Additionally, preventing crosslinking (i.e., stiffening of the collagen fibrils) also reduces the tendon stress response, however, not quite as much as preventing fibril elongation. Furthermore, in contrast to inhibited fibril elongation, loss of crosslinking is predicted to increase the fibril:tissue strain ratio since the softer fibrils will be easier to elongate. In contrast, while the a reduced fibril volume fraction decreases the macroscale stress supported by the tissue, the fibril:tissue strain ratio is nearly unchanged (**Fig. 5C-D**). Additionally, there is virtually no effect of inhibiting the change in fibril diameter on either the macroscale or multiscale response.

**Figure 5:**
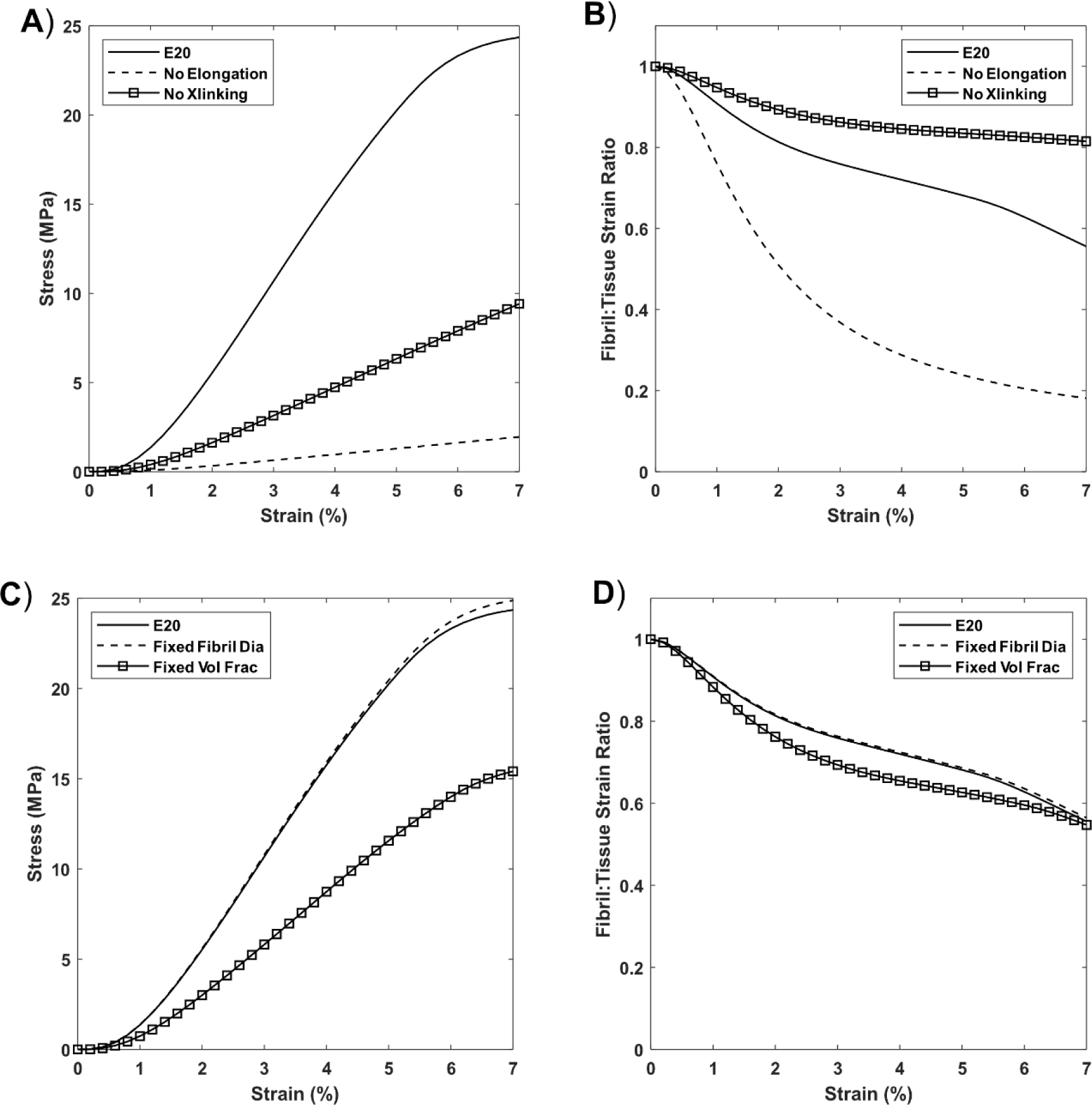
Shear-Lag Simulations of Structural Determinants. **A & B)** Simulated macroscale and microscale mechanical response following the selective inhibition of crosslinking or fibril elongation. **C & D)** Simulated macroscale and microscale response with a fixed fibril diameter or fixed fibril volume fraction.

While it isn’t possible to selectively inhibit fibril elongation or changes in fibril diameter/volume fraction experimentally, we attempted to test the prediction of crosslink inhibition by using a LOX inhibitor. In comparison to saline controls, no significant reduction in embryonic motility was observed following BAPN treatment at E15 (**Fig. 6A**). BAPN treatments resulted in a significant reduction in the macroscale modulus of the tissue in comparison to saline controls (p < 0.001; **Fig. 6B & C**). To investigate how the multiscale mechanics were affected following a reduction of LOX mediated crosslinks, we measured the fibril:tissue strain ratio as a function of applied tissue strain for each group. Both the control and BAPN treated tissue showed an average fibril:tissue strain ratio that was significantly less than one (p < 0.05) (**Fig. 6D**). However, contrary to the elastoplastic model predictions, a linear mixed model found a significant reduction in the fibril:tissue strain ratio following BAPN treatments (p < 0.001) in comparison to saline control.

**Figure 6:**
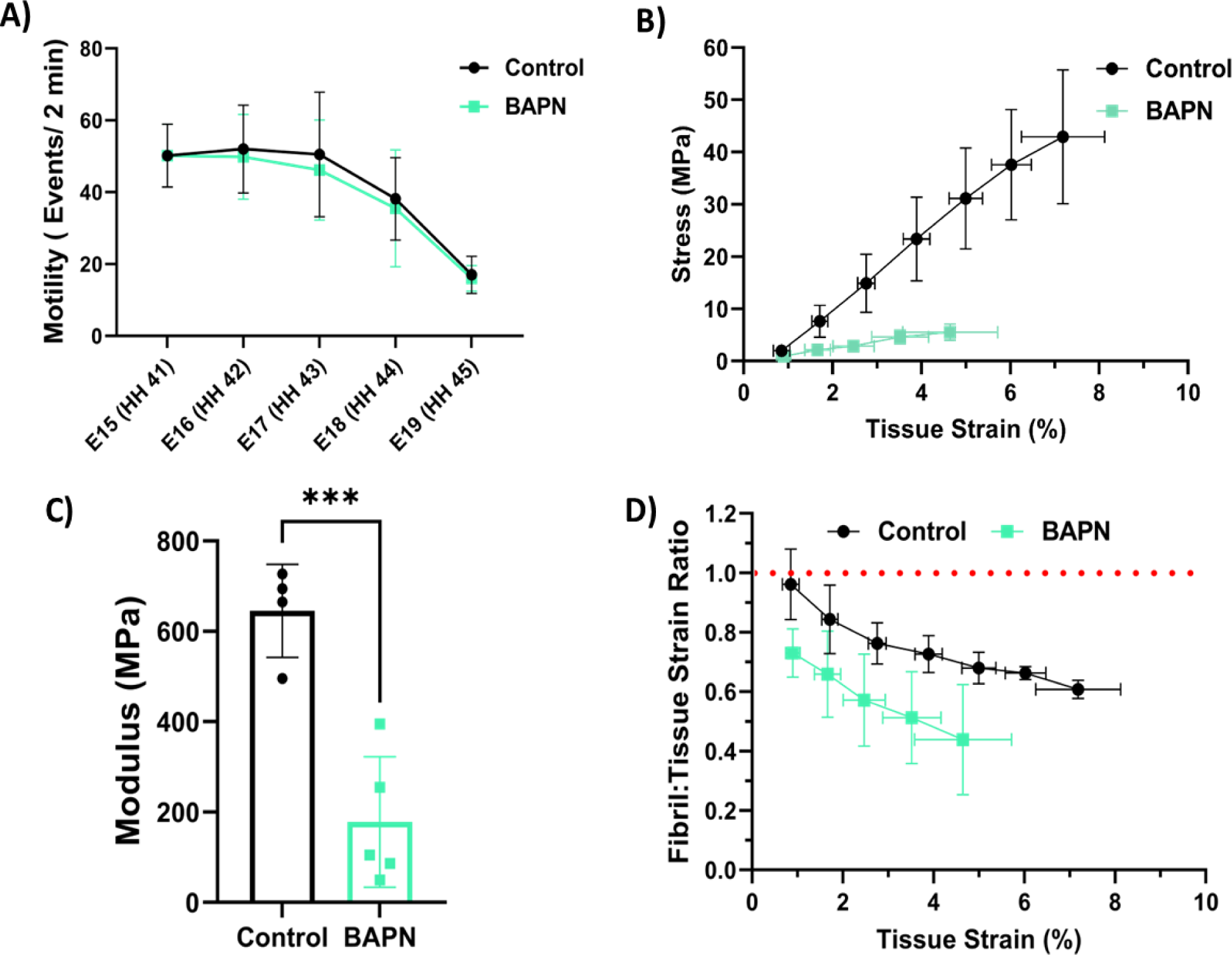
Multiscale mechanical effects following LOX inhibition. A) No significant difference in motility behavior was observed following BAPN treatments. B) Equilibrium stress vs. applied strain for E20 (HH46) samples follow saline (n = 4) or BAPN (n = 5) treatments. C) Equilibrium modulus is significantly reduced following BAPN treatments (p < 0.001). D) Multiscale mechanical response at E20 (HH46) as a function of applied tissue strain after BAPN treatments. The fibril:tissue strain ratio was significantly reduced (p < 0.001) following BAPN treatments in comparison to saline controls. Data represented at mean ± standard deviation. *p< 0.05, ∗∗p < 0.01, p*** < 0.001.

#### Ultrastructural Changes with Crosslinking Inhibition

Following LOX inhibition with BAPN, we observed a significant increase in the spread of the fibril diameters (p < 0.0001) with a particular increase in the number of small diameter fibrils (**Fig. 7**). No significant change in fibril area fraction (p = 0.81) was observed following BAPN treatment in comparison to the vehicle controls.

**Figure 7:**
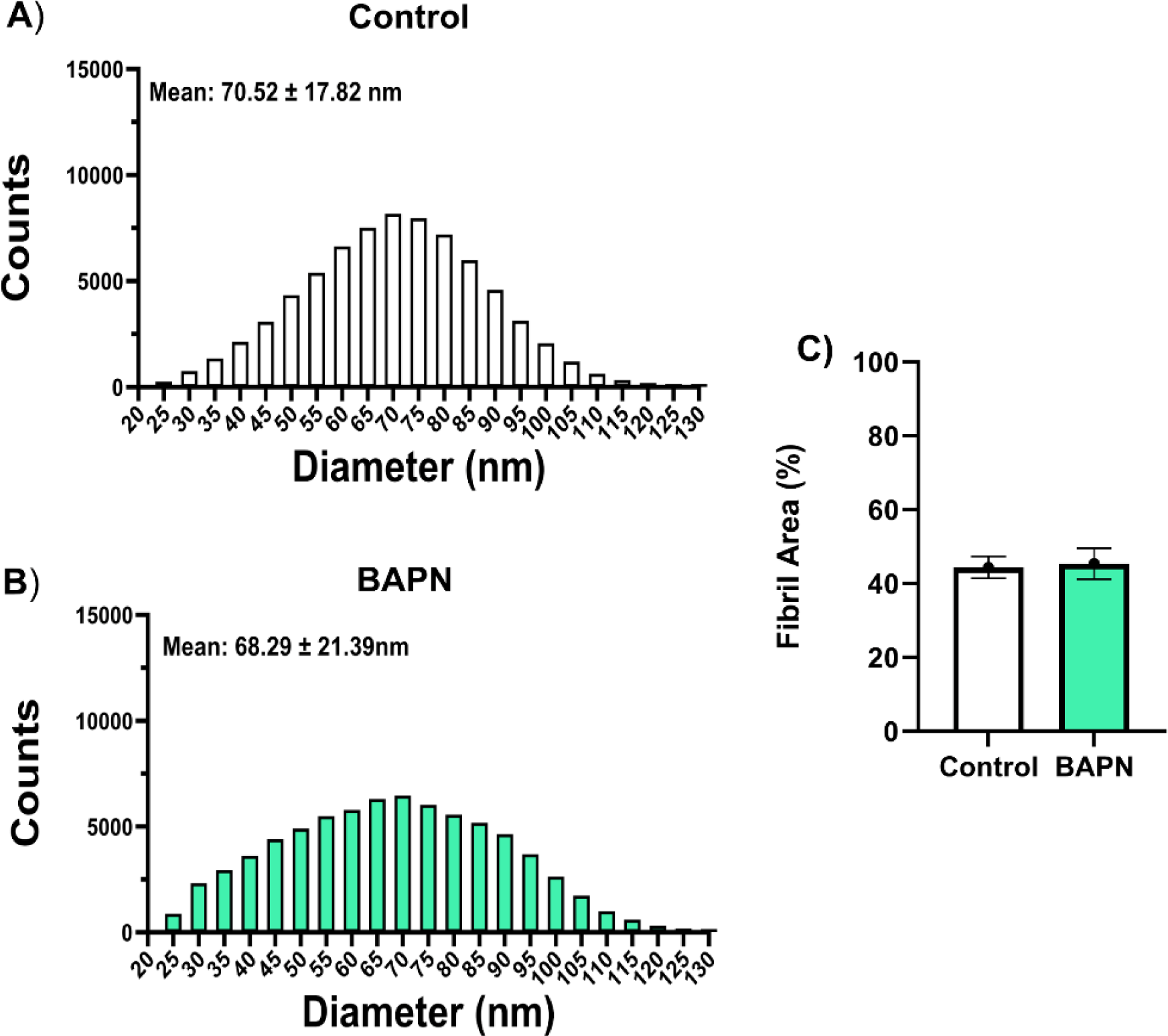
Ultrastructural changes following crosslink inhibition. **A - B)** Fibril diameter distribution following BAPN treatment in comparison to vehicle controls. C) There was no significant change (p = 0.81) in fibril area following BAPN treatment. Each timepoint consisted of two biologicals replicates (n = 2) and six representative regions of interest.

The elastoplastic model fit the control and BAPN treated tissue reasonably well with R^2^ values of at least 0.990 and 0.73 for the stress response and fibril:tissue strain ratio, respectively (**Fig. 8**). Not surprisingly, the model predicted the fibril modulus to be substantially lower with BAPN treatment (**Table 2**). Additionally, consistent with a decreased fibril:tissue strain ratio compared to the saline controls, the model predicted that the fibril length is also substantially smaller with BAPN treatment. In fact, nearly all the model parameters for the BAPN treated samples are very similar to those for the E16 tendons.

**Figure 8:**
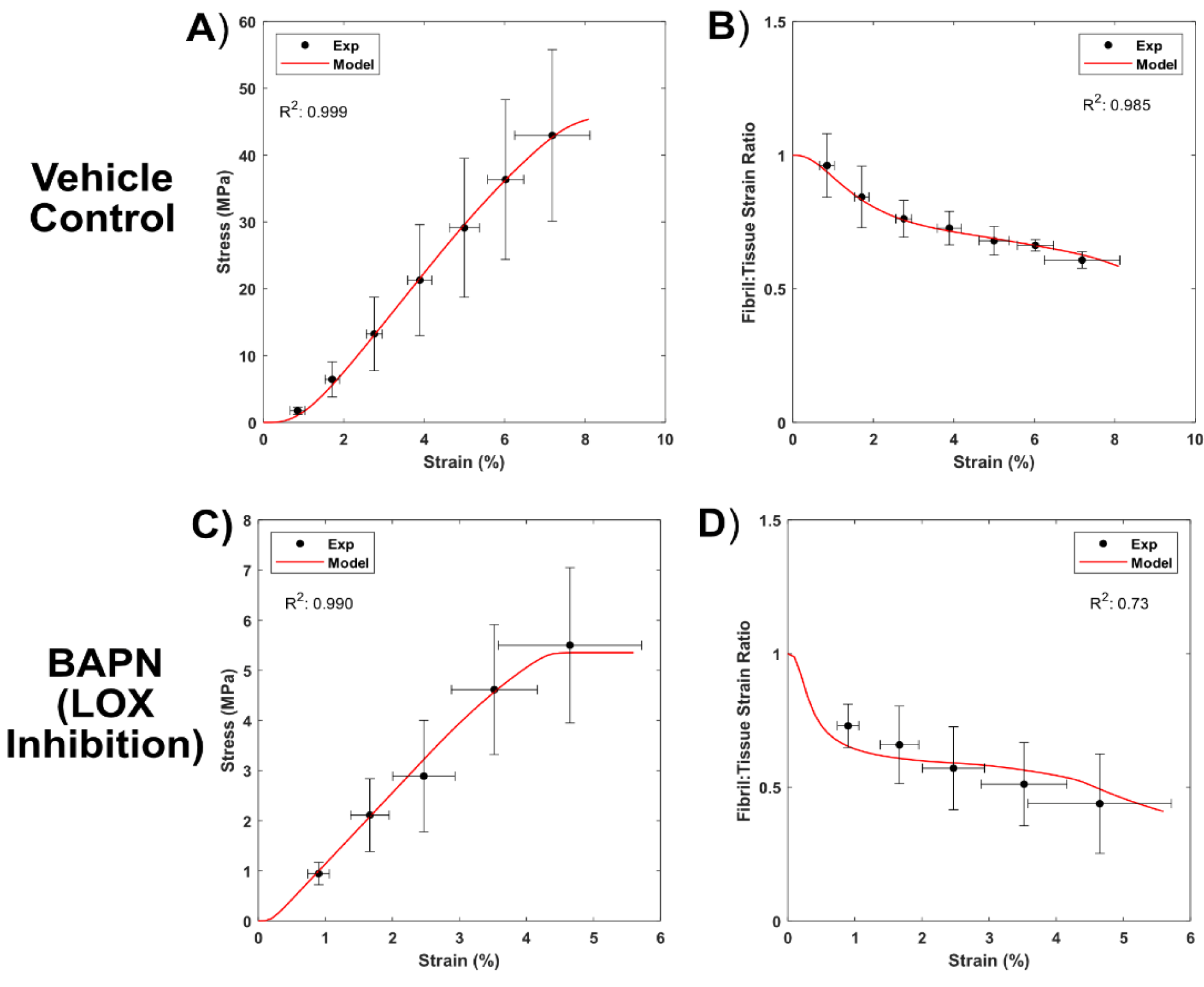
Shear-lag model fit (red line) of the experimental multiscale mechanical behavior (●) following crosslink inhibition at E15 via BAPN. **A & C)** The model successfully fit the equilibrium macroscale mechanics and **B & D**) microscale fibril:tissue strain ratio following the impaired formation of intrafibrillar crosslinks.

**Table2:** Fit shear-lag model parameters following BAPN Treatment (LOX inhibition)

|  | Fibril Modulus (MPa) | Fibril Length (mm) | Avg. Uncrimping Stretch | Std. Dev. Uncrimping Stretch | Shear Modulus (kPa) | Plastic Limit (kPa) |
| --- | --- | --- | --- | --- | --- | --- |
| Control (E20) | 2,792 | 2 (fixed) | 1.0103 | 0.0060 | 0.0086 | 3.7 |
| BAPN (E20) | 565 | 0.22 | 1.0020 | 0.0009 | 0.091 | 3.7 (fixed) |

Following LOX inactivation at E15 with BAPN, we observed a significant reduction (p < 0.05) in total quantity of mature HP/LP crosslinks in comparison to vehicle controls at E20 (**Fig. 9C**). Thermodynamic testing resulted in two distinctive endotherm peaks, but the enthalpy of the Peak 2 population was notably reduced in the BAPN treated samples (p < 0.001) (**Fig. 9D**). The BAPN treated embryos showed no significant difference in the total normalized enthalpy in comparison to their saline controls (**Supplemental Table 2**), indicating LOX inhibition had no effect on the relative abundance of collagen. A reduction in the denaturation temperature (p < 0.01) of the Peak2 population was observed following BAPN treatment. Notably, there was a significant increase in the enthalpy absorbed in the peak 1 (p < 0.001) population relative to the total enthalpy in the system (**Fig. 9E**), characteristic of an abundance of non-crosslinked collagen.

**Figure 9:**
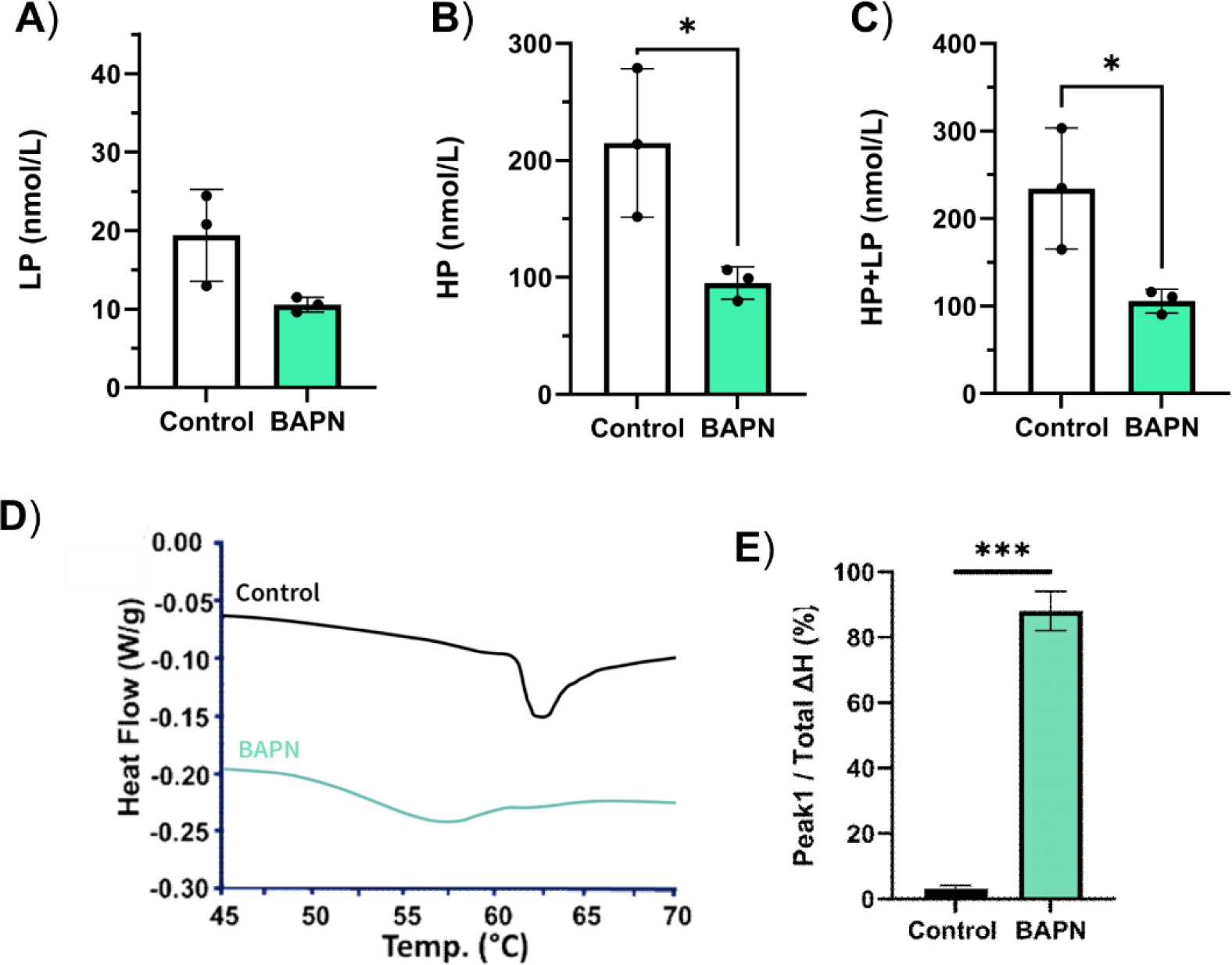
Collagen crosslinking in LOX inhibited samples. Biochemical quantification of mature **A)** LP and **B)** HP crosslinks as a function of crosslink inhibition. **C)** Following LOX inhibition with BAPN (n = 3), there is a significant decrease (p<0.05) in the total mature crosslink populations. **D**) Representative endotherms following crosslink inhibition with BAPN treatment and saline controls. Note that endotherms were shifted along the y-axis to avoid overlap. E) Significant increase in Peak 1 enthalpy / total enthalpy plot following BAPN treatment (p < 0.001). Data represented as mean ± deviation. * p< 0.05 ** p< 0.01, *** p < 0.001

### 3.3 Musculoskeletal Activity is Required for Proper Tendon Development

Rigid (DMB) and flaccid (PB) paralysis treatments significantly reduced embryonic motility through the target period (DMB: p < 0.05, PB p < 0.05) (**Fig. 10A**). No significant difference in motility was observed between the mode of paralysis (p = 0.76). At the macroscale level, both rigid and flaccid paralysis significantly reduced the equilibrium modulus when compared to vehicle control samples (**Fig. 10C**) (p< 0.01 and p < 0.05, respectively). Similarly, a linear mixed model found a significant decrease in the fibril:tissue strain ratio for the both immobilization treatments (DMB & PB) in comparison to saline controls (p < 0.05).

**Figure 10:**
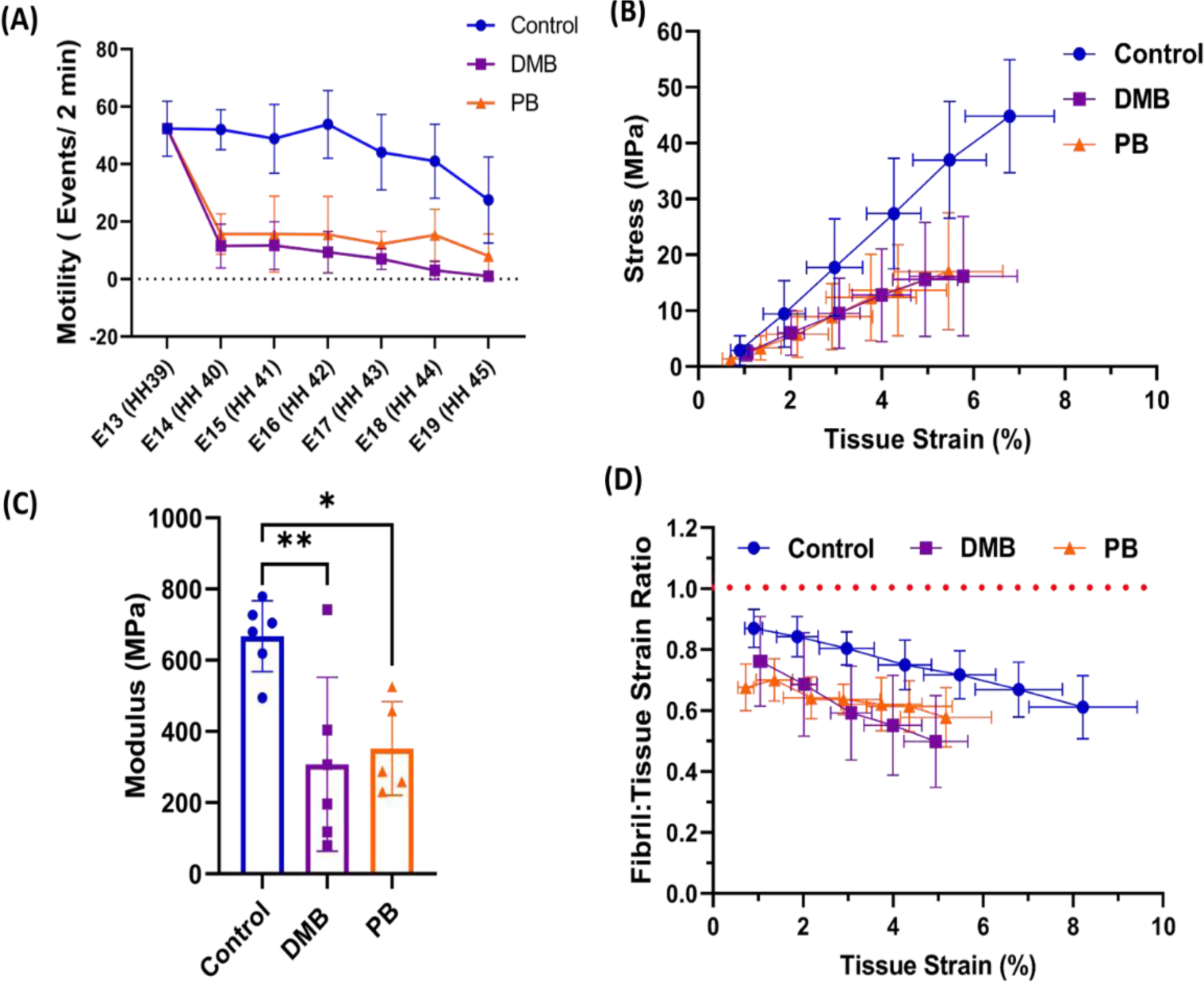
Muscle paralysis and its effect on the multiscale mechanical properties during development. A) Paralysis treatments and saline control were conducted from E13 (HH39) – E19 (HH45). Significant decrease in motility was observed under rigid (DMB, p < 0.01) and flaccid (PB, p < 0.01) paralysis. No significant difference in motility behavior was observed between the two immobilization methods. **B**) Equilibrium stress vs. applied strain for E20 (HH46) samples following saline (n=6), DMB (n = 6), or PB (n= 5) treatments. **C**) Equilibrium modulus is significantly reduced with both rigid paralysis (DMB, p < 0.01) and flaccid (PB, p < 0.05) treatments. **D**) Multiscale mechanical response at E20 (HH46) as a function of tissue strain after immobilization. The fibril:tissue strain ratio was significantly lowered following both rigid (DMB, p < 0.05) and flaccid paralysis (PB, p < 0.05) in comparison to saline controls. Data represented as mean ± standard deviation. ∗p < 0.05, ∗∗p < 0.01.

#### Ultrastructural Changes with Paralysis

Following paralysis treatments, we observed a significant decrease in the number of small diameter fibrils in comparison to vehicle controls (p < 0.0001) (**Fig. 11**). We observed no significant change in fibril area fraction following paralysis treatments, although there was trend toward a decrease for the DMB treated samples (DMB, p = 0.16; PB, p = 0.91).

**Figure 11:**
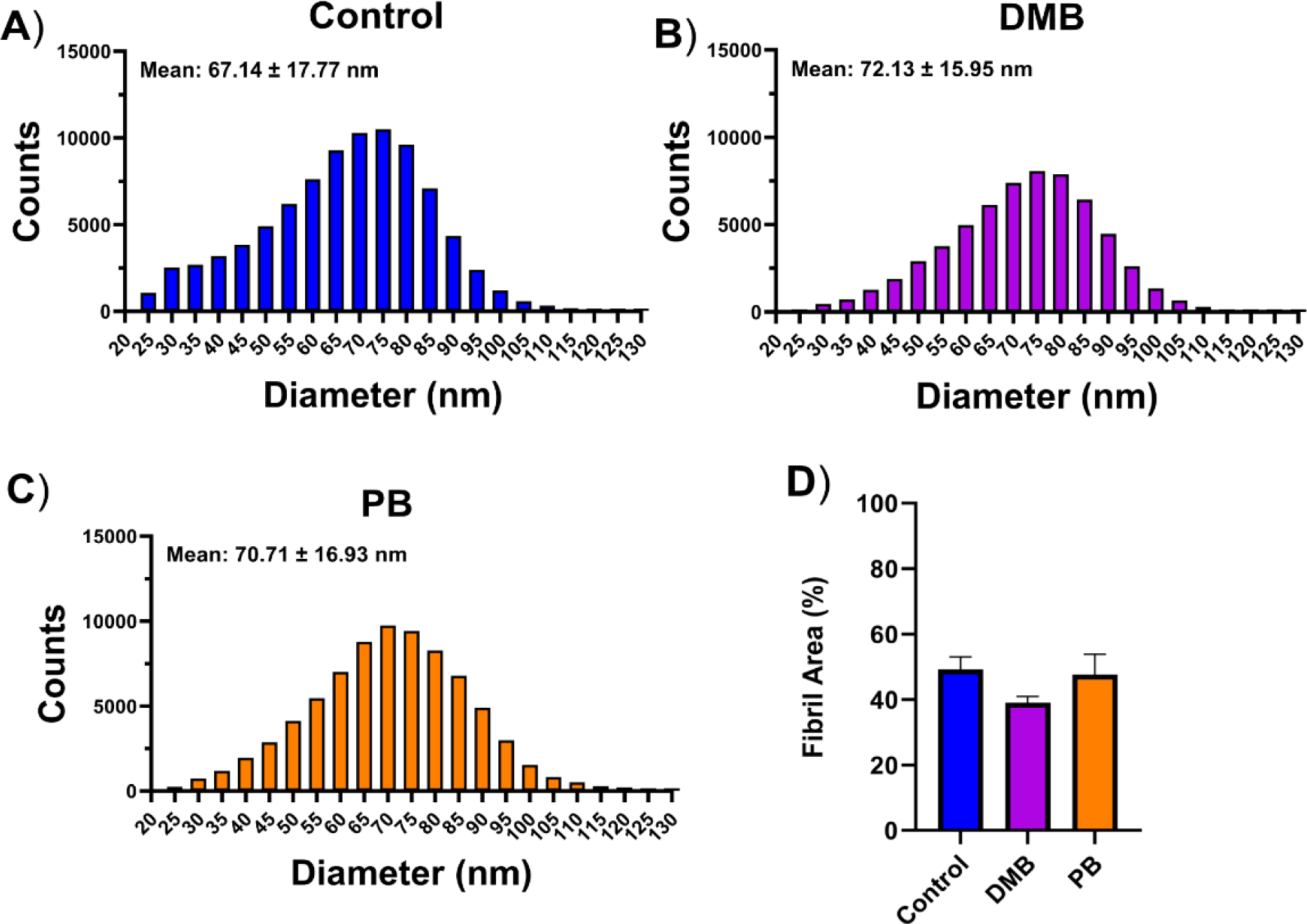
Changes in fibril diameters and area fraction with paralysis. **A - C)** Fibril diameter plots (E20) following rigid (DMB) or flaccid (PB) embryonic paralysis induction at E13. **D)** There was non-significant decrease in fibril area fraction following DMB treatment (p = 0.16) in comparison to saline controls. Each timepoint consisted of two biologicals replicates (n = 2) and six representative regions of interest.

The elastoplastic model was reasonably successful in fitting the paralyzed tendon samples with R^2^ values of 0.995 and 0.90 for the stress response and fibril:tissue strain ratio, respectively (**Fig. 12**). For the PB treated samples, since the fibril:tissue strain data was nearly independent of the applied tissue strain, the R^2^ values of the model fits for these data were poor compared to a horizontal line (−0.94); however, the model still provided a reasonable qualitative approximation of the fibril:tissue strain ratio. The model predicted that both DMB and PB decreased the fibril modulus and the fibril length compared to saline controls (**Table 3**). Additionally, the uncrimping parameters were slightly reduced while the interfibrillar shear modulus was slightly increased, which is similar to tendons at earlier (< E20) developmental timepoints.

**Figure 12:**
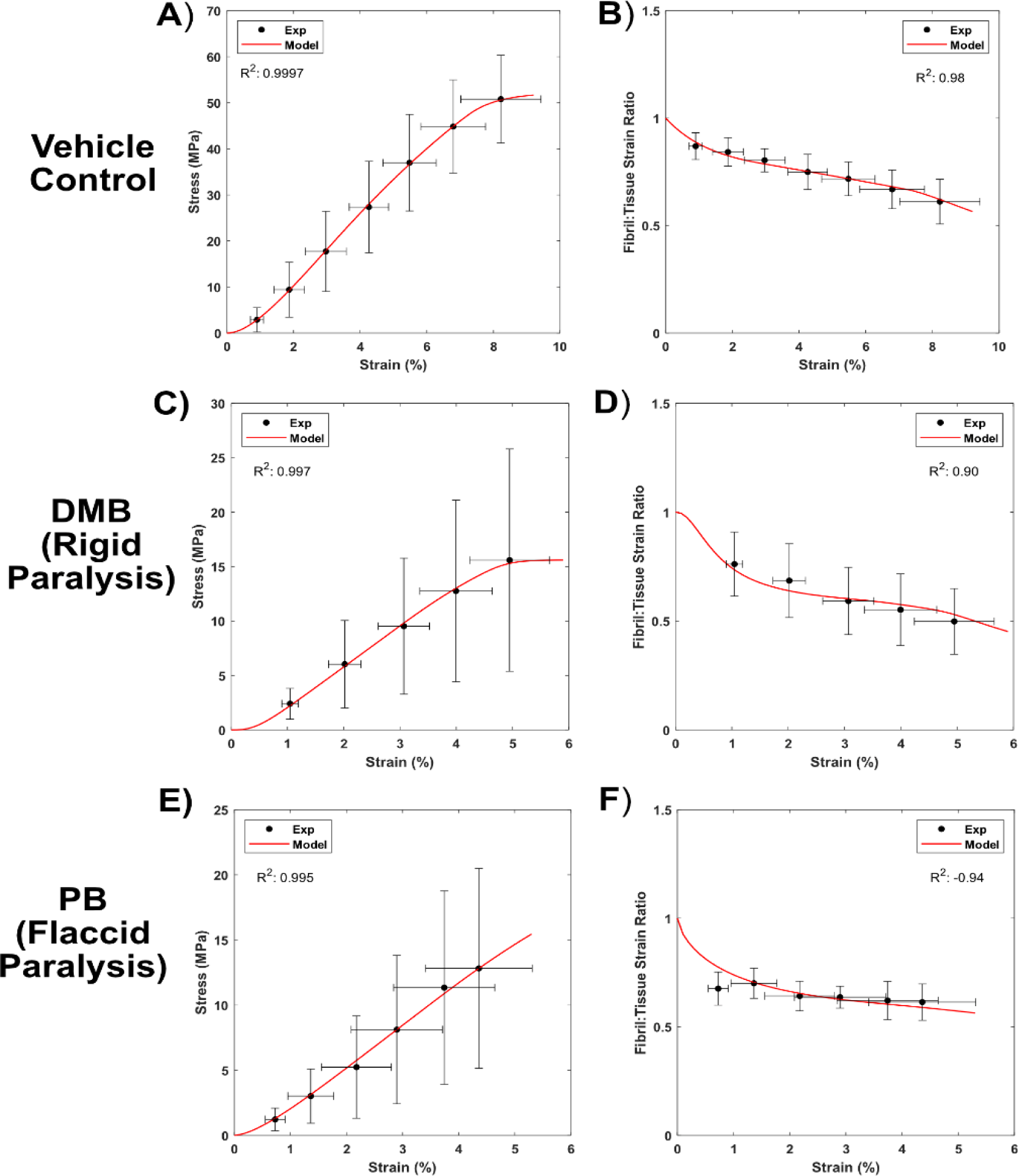
Shear-lag model fit (red line) of the experimental multiscale mechanical behavior (●) following paralysis via DMB or PB treatments at E13. **A, C, & E)** The model successfully fit the equilibrium macroscale mechanics and **B, D, & F**) microscale fibril:tissue strain ratio following embryonic paralysis.

**Table3:** Fit shear-lag model parameters following rigid (DMB) or flaccid (PB) paralysis.

|  | *Fibril Modulus*<br>*(MPa)* | *Fibril Length*<br>*(mm)* | *Avg.*<br>*Uncrimping*<br>*Strech* | *Std. Dev.*<br>*Uncrimping*<br>*Strech* | *Shear Modulus*<br>*(kPa)* | *Plastic Limit*<br>*(kPa)* |
| --- | --- | --- | --- | --- | --- | --- |
| **Control (E20)** | 2,444 | 2 (fixed) | 1.0086 | 0.0086 | 0.011 | 3.6 |
| **DMB (E20)** | 1,826 | 0.8 | 1.0046 | 0.0028 | 0.027 | 3.6 (fixed) |
| **PB (E20)** | 1,336 | 0.88 | 1.0054 | 0.0068 | 0.015 | 3.6 (fixed) |

Following muscle paralysis, there was no significant effect on the presence of total mature crosslinks when measured at E20 (**Fig. 13C**). Furthermore, there was no detectable difference (p = 0.69) between the modes of paralysis (i.e., rigid vs. flaccid paralysis). Following immobilization treatments, neither flaccid (PB) or rigid (DMB) paralysis resulted in a significant reduction in the total enthalpy of the system (p = 0.56 and p = 0.92, respectively) (**Supplemental Table 2**). However, both forms of paralysis resulted in a significant reduction in the denaturation temperature of the Peak 1 and Peak 2 population (p < 0.05) (**Supplemental Table 2)**. Notably, only the flaccid paralysis treatment resulted in a significant increase in the relative enthalpy in the Peak1 population relative to the total enthalpy of the system (Peak 1/ total Δh) (**Fig. 13E**).

**Figure 13:**
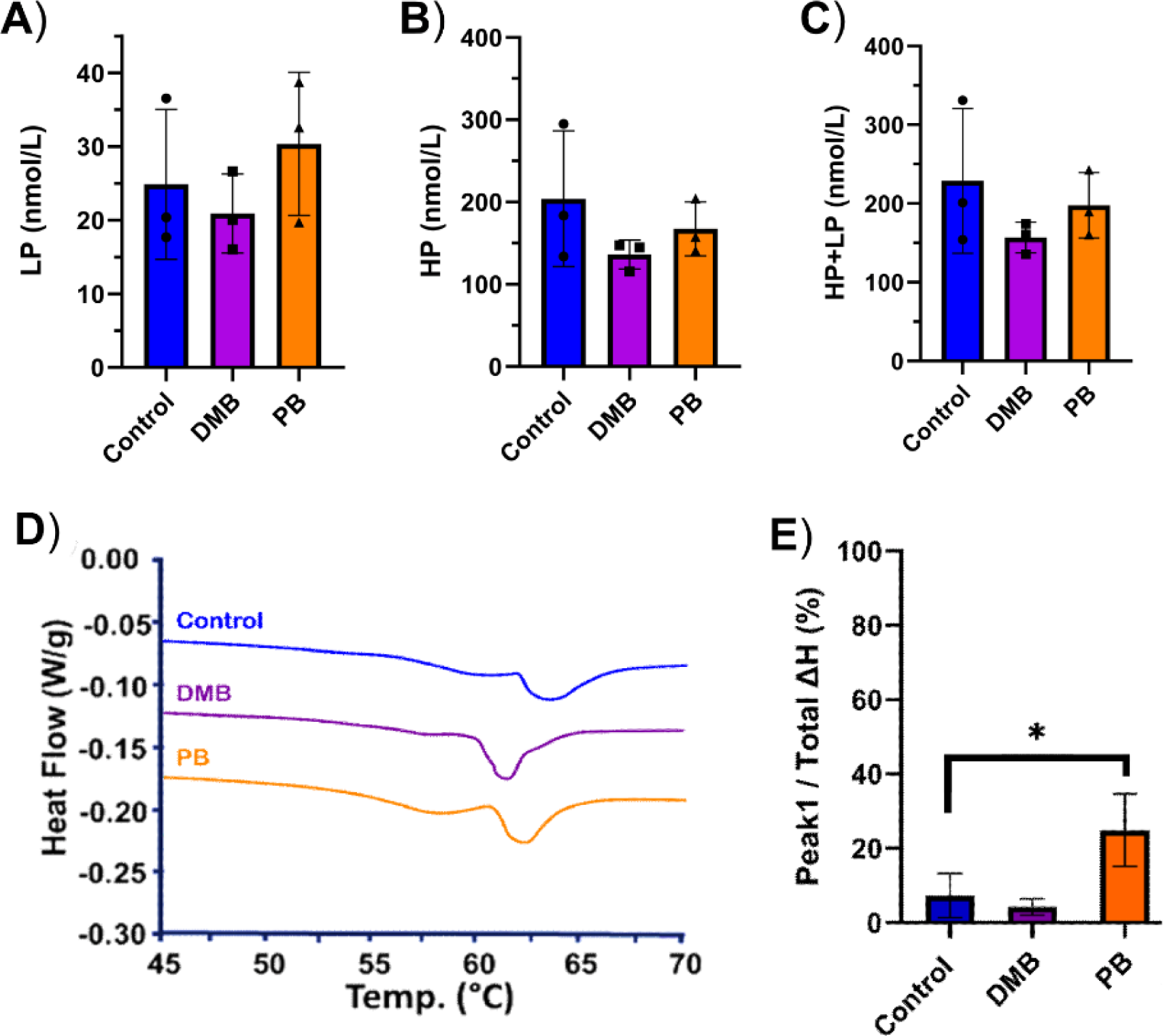
Collagen crosslinking in paralyzed samples. Biochemical quantification of mature **A**) LP and **B**) HP crosslinks as a function of paralysis (n = 3). No significant difference in **C**) HP+LP crosslinks following DMB (p = 0.26) or PB (p = 0.62) paralysis in comparison to vehicle controls. **D**) Representative endotherms of E20 tissue following rigid paralysis (DMB) or flaccid paralysis (PB) in comparison to saline controls. Note that representative endotherm curves were shifted vertically for comparisons across conditions. **E**) A signific increase in Peak 1 enthalpy/ total enthalpy was observed following flaccid paralysis (p < 0.05). Data represented as mean ± standard deviation. * p< 0.05,

## 4 Discussion

This study aimed to elucidate the structural elements that give rise to the load-bearing capabilities of tendon during development and investigate their dependence on muscle activity. Utilizing a combination of mechanical testing, computational modeling, ultrastructural imaging, and crosslinking characterization techniques, we demonstrated that fibril elongation and enzymatic crosslinking are the key players that mediate the structure-function transformation during late stages of development. Changes in volume fraction are also important in terms of the load-bearing capacity but not in terms of the multiscale tissue behavior. Finally, there is minimal importance to the observed changes in fibril diameter. We further demonstrated that muscle activity plays an essential role in mediating fibril elongation during development and that collagen crosslinking is partially mechanosensitive. These findings provide critical insight into the highly dynamic structure-function relationships during tenogenesis and the biological mechanisms in play.

While prior work has postulated that fibril elongation drives load-bearing capabilities in the developing tendon^3,34,35^, directly measuring fibril lengths has proven to be a critical technical barrier. Although collagen fibrils can be extracted and measured from embryonic tendons, intact fibrils cannot be isolated at later developmental stages (>E16 in chicks)^14^. More recent studies have leveraged serial-block face scanning electron microscopy (SBF-SEM) to track fibrils *en bloc*^12,27^, enabling the direct characterization of the collagenous ultrastructure in the native 3D environment. While powerful, SBF-SEM cannot measure fibrils end-to-end as their length scale (∼100 um – 10 mm) greatly surpasses the imaging depth that can be reasonably probed (50 – 100 um), requiring the use of a probabilistic model to estimate fibril lengths^27^. Importantly, this method requires the robust tracking of thousands of individual fibrils to identify distinctive ends throughout the image stack, which has proven to be a large technical challenge.

To overcome these issues, we leveraged a computational shear-lag model^26^ in conjunction with multiscale mechanical testing to estimate the changes in the fibril length and other structural parameters during development. The model demonstrated that the maturation of tendon multiscale mechanics during late stages of chick development (E16 – E20) necessitates a progressive increase in individual fibril length and modulus. More specifically, our model indicated an increase in fibril length from 220 μm to 1,360 μm between E16 and E18, respectively (**Table 1**). Notably, this rapid fibril elongation is consistent with prior observations in embryonic chicks that report an abrupt increase in fibril continuity between E16 and E17^10^. We further saw an estimated 4-fold increase in the fibril modulus with maturation (**Table 1**), consistent with the increase in mature intrafibrillar crosslinks observed with development^36^. The model also indicated that the level of crimping increased with development, which contradicts prior experimental data suggesting that there is actually a small decrease in crimp angle in chick tendons from E16 to post-hatching^37^. However, it is possible that the preload applied prior to mechanical testing extinguished the crimp in the softer E16 tissue, which would have produced a more linear stress response. Finally, the model suggested a novel finding that the interfibrillar matrix becomes more plastic and less elastic with increasing development.

To assess the individual contributions of fibril elongation, collagen crosslink formation, increase in fibril diameters, and increase in fibril volume fraction to the transformation in the load-bearing capabilities of developing tendon, model simulations were performed where each of these structural changes were selectively fixed at their immature state (E16). Notably, while both inhibition of fibril elongation and collagen crosslinking predicted severely impaired bulk mechanical capabilities, they displayed distinctively different multiscale mechanical responses. When fibril elongation mechanisms are selectively impaired, the model predicts a *decrease* in the fibril:tissue strain ratio, indicative of diminished fibrillar strains and increased interfibrillar sliding. This response is consistent with prior work in our lab which has demonstrated a direct link between increased fibril continuity and multiscale mechanical capabilities^24^. Conversely, when intrafibrillar crosslinking (i.e. fibril modulus) is selectively impeded, the model predicted an *increase* in the fibril:tissue strain ratio, indicative of amplified fibrillar stretching. While experimental inhibition of collagen crosslinking using BAPN impaired the macroscale mechanics, we interestingly observed a reduction in the fibril:tissue strain ratio (**Fig. 6**), which is *<u>opposite</u>*to the model predictions and suggests that the fibril elongation may be additionally impaired. Indeed, fitting our shear-lag model to the multiscale mechanical response of the BAPN treated tissue predicts a near 10-fold reduction in fibril length in comparison to vehicle controls, which is even greater than the effect on the fibril modulus. Analysis of the fibril morphology following BAPN treatment showed a decrease in average diameter but a notable widening of the overall distribution (**Fig. 7B**), suggesting that LOX may mediate fibril interactions during development. Indeed, BAPN treatments on *in vitro* tendon constructs reported irregular fibril morphologies^18^. While the formation of enzymatic crosslinks is well understood to contribute to the mechanical capabilities of tendons^15,38,39^, it’s unclear if LOX interacts with other collagens (i.e., V or XI) or proteoglycans (i.e., decorin, biglycan, lumincan, or fibromoduluin), which have been implicated in mediating fibril interactions^40–47^. All together, these finding suggest that LOX-mediated crosslinking plays a key role in the establishment of the structure-function relationships during development, but additional work needs to be done to identify the mechanisms by which LOX influences fibrillogenesis.

Having established the structure-function relationships during development and following LOX inhibition, we then aimed to better understand the role of muscle activity in tenogenesis. Following the induction of paralysis at E13, we saw a significant reduction in the multiscale mechanical capabilities (**Fig. 10**) regardless of the form paralysis. Interestingly, we observed a small, but highly significant (p< 0.0001) increase in fibril diameter following either form of paralysis (**Fig. 11**). These data suggest that lateral fibril interactions were not impeded in the absence of muscle stimulation. Intrestingly, biochemical assays reported no significant difference in the total quantity of mature crosslinks following paralysis (**Fig. 13C**). However, thermodynamic measurements show noticeable effects on the thermal properties following flaccid paralysis. More specifically, there is a reduction in the denaturation temperature (Supplemental Table 2) and an increase in the Peak1/total enthalpy measurements in comparison to the saline control. The discrepancy between the DSC and biochemical data could be due to immature divalent crosslinks that are not detectable in the biochemical assays or because other factors (e.g., tissue hydration) may be contributing to the thermal stability of the tissue. When the shear-lag model is fit to the multiscale mechanical response, we estimated a decrease in fibril length and modulus for both treatments; however, flaccid paralysis produced a larger decrease in fibril modulus, matching our thermodynamic assessments. Together, these data suggest that the mechanosensitivity of collagen crosslinking is dependent on the mode of paralysis, with flaccid paralysis having a more pronounced effect. This may be explained by the fact that DMB induces an irreversible muscle contraction, generating a prolonged static load across the tissue^16^, whereas PB inhibits muscle contraction via competitive binding of acetylcholine at the motor plate, causing a reduction in muscle function altogether^16^.

While this study has provided some critical insight into the dynamic structure-function capabilities during development, there are a few notable limitations. First, while the utilization of the shear-lag model providing insight into the structure-function relationships of developing tendon, we were unable to experimentally validate the predictions. That is, we were unable to directly measure fibril lengths at various developmental stages or treatments. We did attempt to measure the fibril lengths using SBF-SEM. However, it was impossible to confidently identify the fibril ends. Therefore, we were limited to standard 2D measurements of fibril diameters and area fraction. While our fibril area fraction data had limited biological replicates, our trends observed with development are consistent with previous reports in the field^2^. It is also important to note that we did not possess the ability to measure individual fibril mechanics to confirm individual fibril modulus estimates. Furthermore, in order to properly constrain the model, we needed to artificially fix a value for the plastic limit of the interfibrillar shear stress across developmental ages and treatments. Despite this, our predicted structural and functional changes generally agreed with our biochemical and thermodynamic assays, as well as prior work in the field. It’s additionally important to note that while BAPN is a well-known inhibitor of the LOX family of enzymes, it is possible that BAPN is interacting with unknown targets. It’s additionally important to note that while BAPN didn’t impair embryonic motility, there was notable morphological phenotype of the developing hindlimb. More specifically, there was impaired fusion of the ankle joint and the calcaneal tendon was improperly positioned. While limbs were still moving during development, it’s unclear how this skeletal morphology affected mechanobiological stimuli during maturation.

Overall, our findings indicate that mechanisms of fibril elongation and crosslink formation are the dominate structural elements that enable the maturation of tendon and its multiscale mechanical capabilities. Furthermore, both of these structural changes are mechanosensitive. This study has provided essential insight into the structural elements which give rise to the load-bearing capabilities during tendon development and their dependence on external mechanical stimulation. Future work aims to identify the biological mechanisms driving these structural changes and their resulting effects on tissue mechanics. This information is important for creating potential opportunities for advancing tissue engineering of tendons.

## 6 Supplemental Figures

**Supplemental Table 1:** Initial Inputs for the Elastoplastic Model Fitting of E20 Controls.

| *Model Parameter* | *Value* | *Justification* |
| --- | --- | --- |
| *Fibril Modulus ($E_f$)* | $E_f = 1.5 \frac{E}{\phi}$ | Fibril modulus is greater than macroscale modulus (E) due to fibril volume fraction ($\phi$) and crimping |
| *Fibril Length ($L$)* | 2 mm (fixed) | Kalson eLife paper |
| *Avg Uncrimping Stretch* | 1.01 | Qualitative inspection of stress-strain curves for initial 2% strain increment (Fig. SX) |
| *Std. Dev. of Uncrimping Stretch* | 0.002 | Szczesny & Elliot, 2014 <sup>29</sup> |
| *Elastic Interfibrillar Shear Modulus* | 0.5 Pa | Szczesny & Elliot, 2014 <sup>29</sup> |
| *Plastic Limit of Interfibrillar Shear Stress ($\tau$)* | $\tau = \frac{4rE_fGL\epsilon_{max}}{8rE_fh + 3GL^2}$ | Value chosen such that macroscale plasticity occurs at tissue failure ($\epsilon_{max}$) |

*Note: $r$ is the fibril radius determined from ultrastructural imaging and $h$ is the surface-to-surface distance between fibrils, which was calculated assuming hexagonal packing (cite).*

**Supplemental Table2:** Thermal properties of embryonic tendons as a function of development or paralysis/crosslink inhibition. Statistical comparisons for development are done with respect to E20 timepoint and treatments are compared to their respective vehicle controls.

| | $\Delta h$ (J/g) | | $T_{peak}$ (°C) | |
| --- | --- | --- | --- | --- |
| | Total $\Delta h$ | Peak 1/ Total $\Delta h$ (%) | Peak1 | Peak2 |
| **E20** | $2.56 \pm 0.46$ | $6.39 \pm 4.95$ | $59.67 \pm 0.33$ | $63.05 \pm 0.57$ |
| **E18** | $1.97 \pm 0.22$ | $20.86 \pm 2.0^a$ | $58.47 \pm 0.23^a$ | $62.29 \pm 0.30^a$ |
| **E16** | $0.88 \pm 0.18^b$ | $16.4 \pm 7.97$ | $57.02 \pm 0.15^b$ | $61.46 \pm 0.22^b$ |
| **Control (E20)** | $2.53 \pm 0.43$ | $3.11 \pm 1.03$ | $57.93 \pm 0.95$ | $63.60 \pm 0.25$ |
| **BAPN (E20)** | $2.02 \pm 0.15$ | $89.14 \pm 7.95^b$ | $57.13 \pm 0.72$ | $62.35 \pm 0.71^b$ |
| **Control (E20)** | $2.06 \pm 0.47$ | $7.28 \pm 5.99$ | $59.46 \pm 0.38$ | $63.18 \pm 0.46$ |
| **DMB (E20)** | $1.92 \pm 0.59$ | $4.25 \pm 2.18$ | $57.61 \pm 0.47^b$ | $61.74 \pm 0.36^b$ |
| **PB (E20)** | $1.67 \pm 0.06$ | $24.87 \pm 9.76^b$ | $57.77 \pm 0.17^b$ | $62.04 \pm 0.25^a$ |

<sup>a</sup> Statistically Significant ( $p < 0.05$ )
<sup>b</sup> Statistically Significant ( $p < 0.01$ )

**Supplemental Figure 1:**
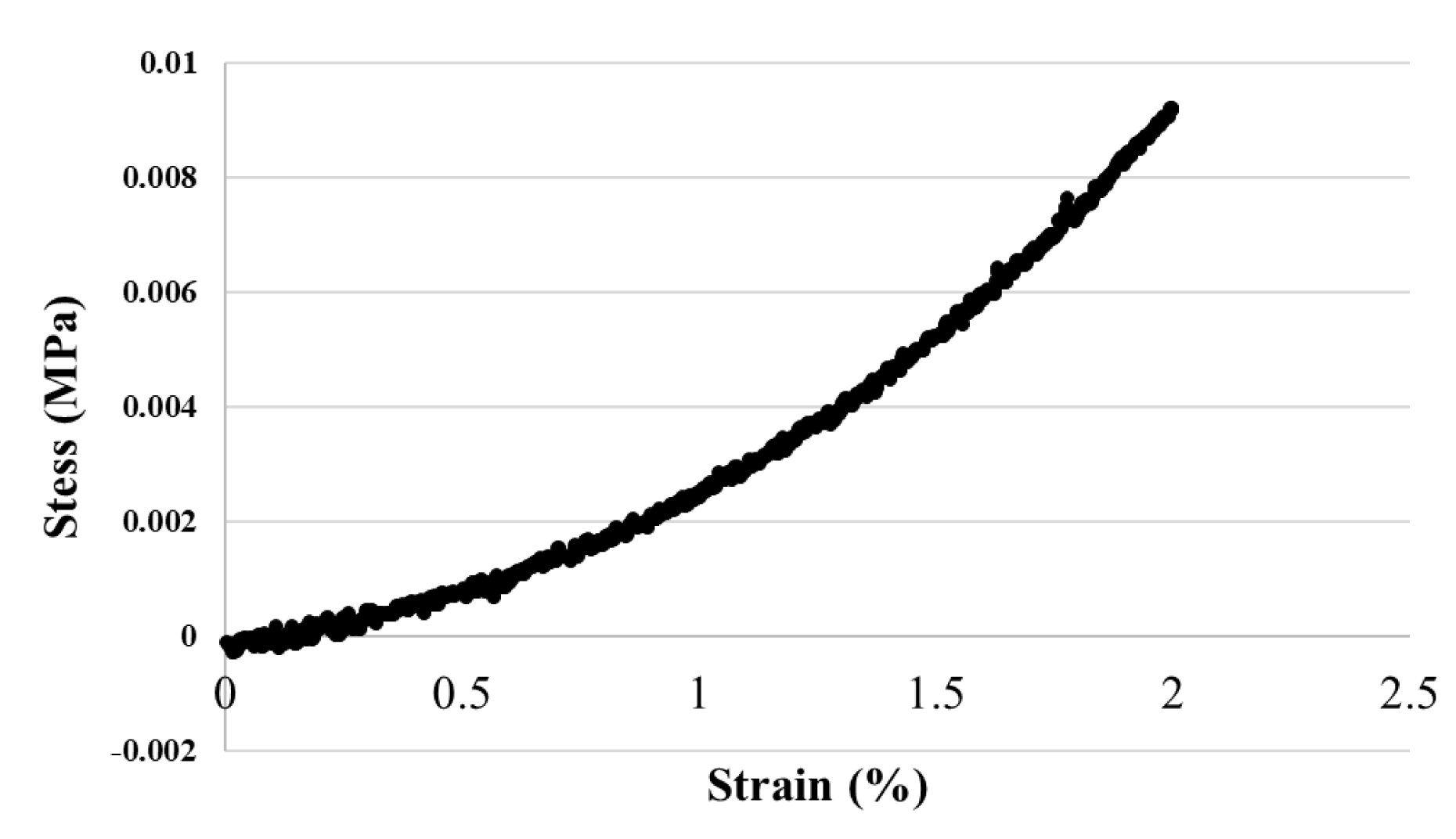
Representative stress-strain curve for E20 tissue.

## Notes

### Competing Interest Statement

The authors have declared no competing interest.

